# Automated Cell Lineage Reconstruction using Label-Free 4D Microscopy

**DOI:** 10.1101/2024.01.20.576449

**Authors:** Matthew Waliman, Ryan L Johnson, Gunalan Natesan, Shiqin Tan, Anthony Santella, Ray L Hong, Pavak K Shah

## Abstract

Here we describe embGAN, a deep learning pipeline that addresses the challenge of automated cell detection and tracking in label-free 3D time lapse imaging. embGAN requires no manual data annotation for training, learns robust detections that exhibits a high degree of scale invariance and generalizes well to images acquired in multiple labs on multiple instruments.

## Main

The study of development is inextricably linked to the concept of lineage at multiple levels.Notable recent advances have led to the development of highly accurate and scalable automated pipelines for reconstructing cell lineages from fluorescence imaging^1–4^. Despite this, manual lineage tracing in label-free images continues to play an important role in the field, ever since its use in seminal lineage tracing experiments of developing embryos^5^. Label-free imaging for small organisms is typically more accessible than high resolution long-term fluorescence imaging, removing the need for transgenesis or staining. Automation, however, has been essential for processing datasets large enough to study variability and robustness. Using stable transgenic reporter lines and well-optimized imaging protocols have facilitated the analysis of several thousands of individual embryos in a variety of contexts^6–10^.

Image processing approaches have not been extensively developed to detect and track cells in widely used 3D label-free imaging modalities such as differential interference contrast (DIC). A range of deep neural network-based approaches have recently been developed to manipulate label-free images including computational staining, where the network is trained to identify subcellular structures in label-free images, and style transfer, where networks transform the appearance of images between modalities^11–13^. These approaches use paired labeled and label-free images of stained or fluorescent samples, minimizing the laborious manual generation of training data typically associated with deep learning approaches. These methods have widely used training schemes that minimize a pixel-by-pixel measurements of intensity mismatch between the network’s output and the real fluorescence modality images provided in the training set. Learning style transfer using a pixel intensity-based loss function attempts to produce a transformed image that closely resembles the target image. For cell detection and lineage tracing, however, visual similarity is less important than the processed image’s compatibility with object detection algorithms.

Reasoning that an object-focused loss function that incorporates a segmentation metric would allow a model to better learn the cell detection task, we used a strategy originally developed for segmenting partially obscured objects in natural scenes^14^ in embGAN, our pipeline for cell detection and tracking in 3D DIC images (Figure 1A). In embGAN a U-Net^15^ generates an object probability map of the DIC image. We incorporated a multi-scale objective loss function^16^ that is computed using the output of a second network functioning as a critic of the segmentation. The critic uses imperfect labels generated by a pre-trained 2D segmentation model^17^ and compares them against the segmentor’s output. This strategy allows both the segmentor and critic to learn both object and image properties, ensuring more robust generalization compared to pixel-wise image mismatch-based loss functions. In our experience, common adversarial training approaches (ie. pix2pix or cycleGAN) using only fluorescence images and the output of the segmentor as inputs to the critic frequently produces unstable networks that fail to converge or generate reliable predictions on images outside the training set. This brittleness is a well-known challenge with adversarial training^18^, which we addressed by segmenting the fluorescence channel to mask cell nuclei, focusing the attention of the segmentor on object detection performance. We found that even imperfect segmentations by a generalist model dramatically improved the performance and robustness of adversarial training in this task.

**Figure 1.**
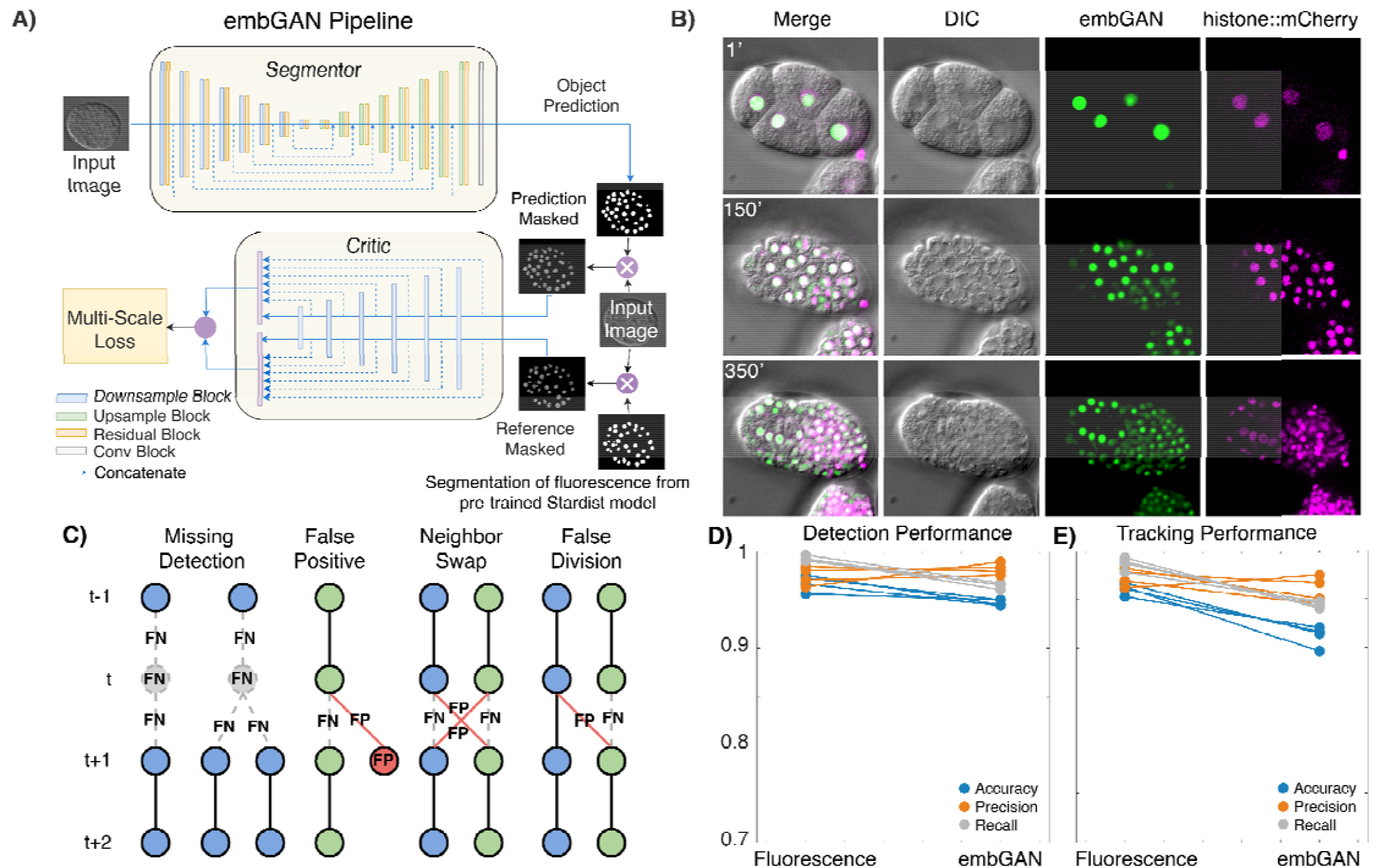
A) Schematic representation of the embGAN network architecture. The Segmentor network generates a segmentation mask of the input image, which is supplied, along with a segmentation generated from the corresponding fluorescence image by a pre-trained Stardist model, to the Critic network whose output is used to calculate the multi-scale loss which is minimized to train the Segmentor and maximized to train the Critic. B) Example 2D images from the validation image set of a *C. elegans* embryo showing the embGAN probability score (green) along with the input DIC image (gray) and nuclear-localized mCherry (magenta) channel. C) Schematics of common detection and tracking errors. Circles represent cell detections; gray circles show FN and red circles show FP. Lines show tracked edges between detections across time points. Dashed lines show FN edges and red lines show FP edges. D) Detection and E) Tracking performance measured on 4 embryos. Each point shows the measurement of Accuracy (blue), Precision (orange), and Recall (gray). Lines connect performance measurements between StarryNite performance on the same embryo’s fluorescence imaging (left) or embGAN-processed DIC imaging (right).

We tested embGAN on a lineage tracing task with a well-characterized ground truth: reconstructing the embryonic lineage of the nematode *Caenorhabditis elegans*^5^. We acquired training data using time lapse 3D microscopy, imaging embryos in both label-free DIC and fluorescence to capture a ubiquitously expressed nuclear-localized mCherry marker. We measured real-world performance using additional time lapse images that were acquired separately from and not included in the training set. Using StarryNite^3^, a well-characterized and robust cell tracking pipeline, we separately analyzed the fluorescence time lapse and DIC images processed by embGAN for 4 embryos (Figure 1B). We then curated both sets of results from all 4 embryos for the first 200 minutes of development (a total of 1,600 volumes up to approximately the 300-cell stage) and compared StarryNite’s output to the curated results. We counted all detection (Tables S1, S2) and tracking (Tables S3, S4) errors (Figure 1C) in each dataset. StarryNite performs extremely well on cell detection tasks with fluorescence images (Figure 1D), with recall exceeding 99% (99% of all cells are detected) and precision averaging 97.4% (2.6% of detections are false positives). For tracking, StarryNite performance on fluorescence images achieved recall and precision averaging 99% and 95.9% (Figure 1E), respectively. Since embGAN is trained using a critic fed with imperfect segmentations, StarryNite cell detection and tracking performance drops slightly compared to corresponding fluorescence images. Despite this, embGAN performance averaged 98.2% recall and 96.4% precision for detection (Figure 1D) and averaged 95.9% recall and 94.4% precision for tracking (Figure 1E).

Since embGAN performed well for lineage reconstruction in the early embryo, we next tested embGAN performance at later stages of development when cells are much smaller and more densely packed. We used the 4 test embryos to identify all 24 neurons of the amphid sensory organs that are born in the final 9th or 10th generations of the AB lineage (Figure 2A,B). Eight of these neurons are born in the 10th generation, which sometimes occurs after the beginning of rapid embryo movement. We tracked these neurons to the birth of their parents in the 9th generation since embryo movement currently prevents reliable tracking. We counted tracking errors in the lineage history of these cells, finding that most errors occur due to cell divisions, either the cell’s own or a nearby cell’s, such that a similar proportion of errors occurred in each generation of the AB lineage. Since the length of the cell cycle of each generation also increases, this results in a decrease in the per-frame error rate at later generations compared to earlier generations starting at 3% in the early embryo and dropping to approximately 1% (Figure 2C).

**Figure 2.**
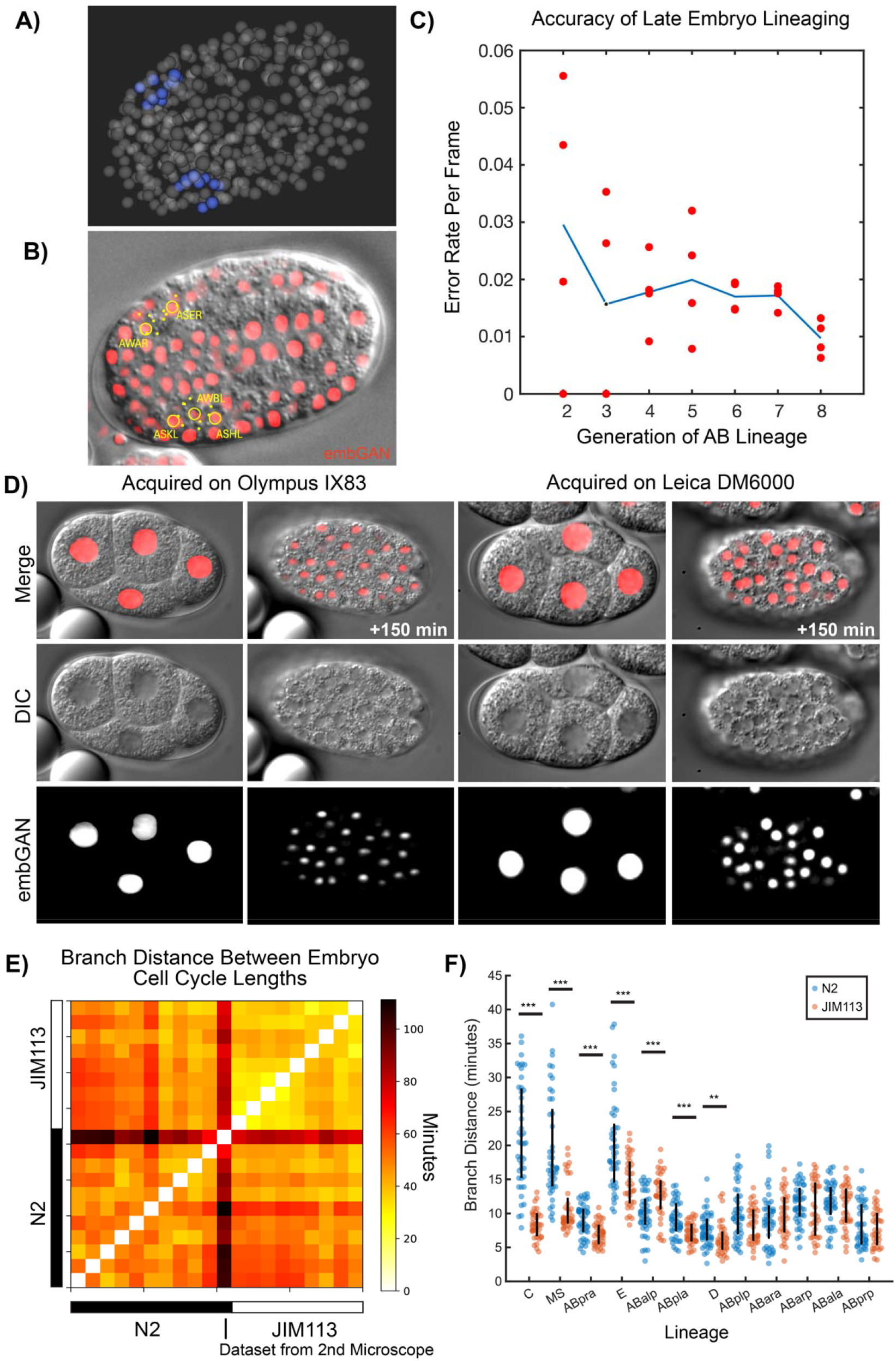
A) 3D rendering of amphid cell positions traced using embGAN images B) Image of DIC and embGAN (red) image with annotations showing positions of amphid cells. Cells present at the focal plane are circled with dots showing the XY position of cells present at different depths in the volume.C) Accuracy of lineaging cells born late in embryogenesis. Mean (blue line) and per-embryo (red circles) error rates for lineaging amphid neurons in embGAN processed images. D) Generalizability of embGAN model across multiple source microscopes. Images of early and mid-stage N2 embryos shown in DIC, embGAN output, and merged channels acquired on the same microscope as the embGAN training data (Olympus IX83) and a second microscope using different hardware (Leica DM6000). E) Heatmap showing the pair-wise branch distance comparing lineage-aligned cell cycle lengths between all N2 and JIM113 embryos. The single N2 embryo imaged using the 2nd microscope is highlighted at the boundary between N2 and JIM113 embryos. F) Distribution of pairwise branch distances between specific lineages within N2 (blue) and JIM113 (orange) embryos. Vertical black line shows the interquartile range (the 25th to the 75th percentile). ^***^ denotes p<0.001 and ^**^ denotes p<0.01 by the rank sum test.

We next tested embGAN’s ability generalize to images of animals from different strains and to images acquired on a different microscope than the training set. For this we imaged the laboratory-raised wild-type strain N2 and used embGAN to reconstruct the early embryo. We imaged N2 embryos using the two different brands of microscope (Leica DM6000 vs Olympus IX83), different objectives and immersion media (63x oil vs 60x silicone oil immersion), and different cameras (Leica K5 vs Hamamatsu C14440-20UP). Using embGAN and StarryNite, we then traced embryonic development through the 6th round of division of the AB blastomere and all other sub-lineages of the embryo to a similar time-point in the 10 N2 embryos imaged using the Olympus microscope and one of the N2 embryos imaged using the Leica microscope (Figure 2D). We used the branch distance, a metric we previously developed to compare lineage-aligned phenotypic measurements between cell lineages^19^, to compare the lineage-aligned distribution of cell cycle durations between the same cells in N2 and a set of transgenic JIM113 embryos (Figure 2E). The embryo imaged using the Leica microscope was a clear outlier here due to a difference in ambient temperature during imaging (approximately 23 □vs 21□) causing a change in the global rate of development (Figure S1).

Interestingly, N2 embryos exhibited ∽2× more inter-embryo variability in the branch distance than JIM113 embryos. This increased variability reflects differences in cell cycle timing between N2 embryos and originates consistently from specific lineages. N2 embryos are more heterogeneous than N2 among all lineages with significant inter-strain differences in the median branch distance between embryos except for one-ABalp (Figure 2F). Whether strain-specific patterns of inter-individual variability originate from the transformation and selection steps in *C. elegans* transgenesis or is common among lines sourced from distinct wild populations is an interesting question that could be addressed using our approach.

Automated pipelines for cell detection and tracking in 3D time lapse fluorescence microscopy have advanced by leaps and bounds in recent years. These efforts have principally focused on improving performance on image sets of organisms too large to generate large amounts of manual annotation for deep learning^1,2^. Label-free imaging remains an attractive technology for the study of early development as doesn’t require transgenic or stained samples. The approach we employ in embGAN makes it possible to achieve the performance required for automated lineage tracing using label-free images. Using higher information content imaging modalities such as quantitative phase imaging (QPI), larger and more diverse training sets, or higher performance fluorescence segmentation models for training data annotation will likely further improve embGAN performance. embGAN makes high throughput cell lineage analysis possible for a wider range of sample types than was previously possible. To accelerate continued improvements in this area, we are making our full training image set freely available (https://datadryad.org, DOI: 10.5061/dryad.zcrjdfnkz) alongside the codebase for embGAN (https://github.com/shahlab-ucla/embGAN; DOI: doi/10.5281/zenodo.10535870).

**Figure S1.**
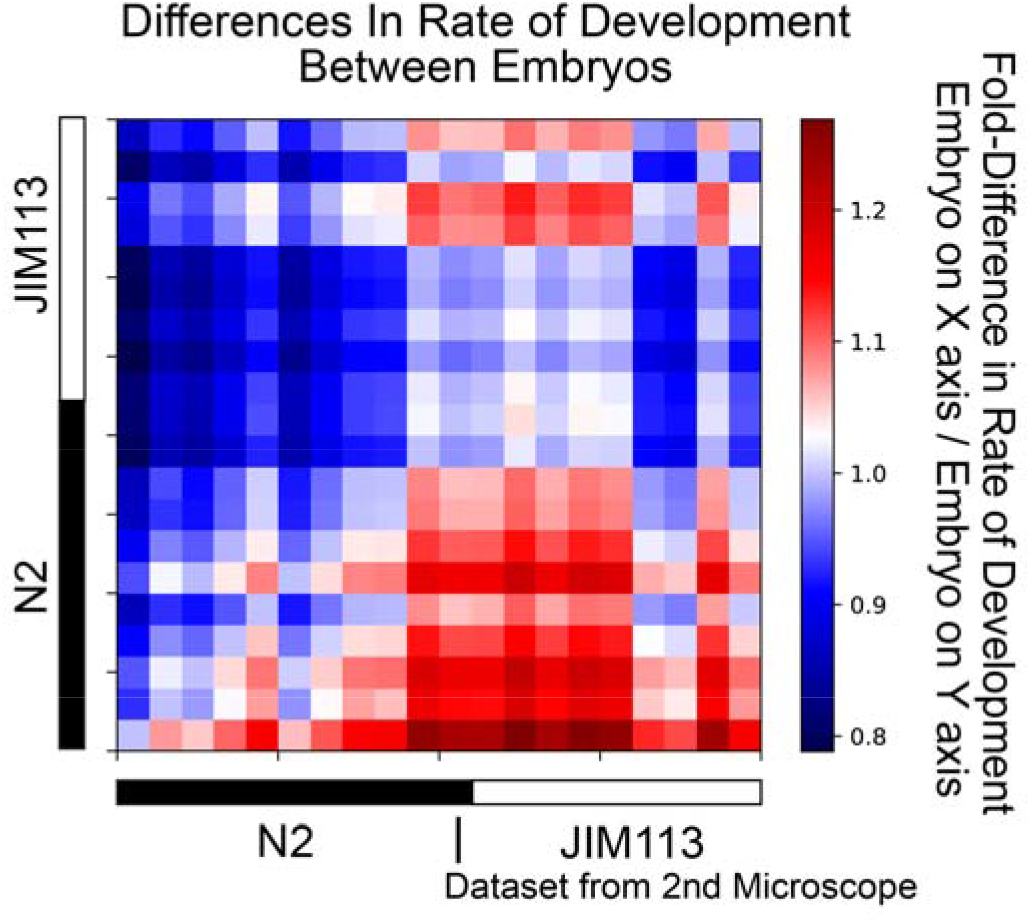
Comparison of global rate of development between all N2 (black bar) and JIM113 (white bar) embryos calculated pairwise using the first principal component as a non-parametric estimate of scaling between embryos. Each square represents a single pairwise comparison where the color reflects the fold-difference in cell cycle durations of the embryo indicated by the x-coordinate of the square relative to the embryo indicated by the y-coordinate of the square.

**Supplemental Table 1.**
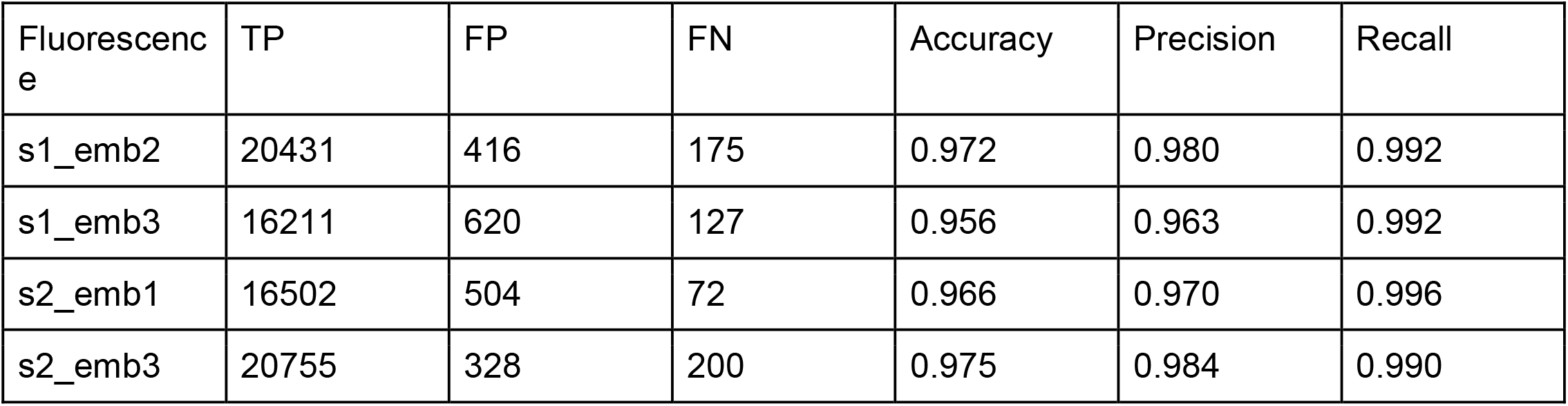
Cell Detection Performance on Fluorescence Images.

| Fluorescence | TP | FP | FN | Accuracy | Precision | Recall |
| --- | --- | --- | --- | --- | --- | --- |
| s1_emb2 | 20431 | 416 | 175 | 0.972 | 0.980 | 0.992 |
| s1_emb3 | 16211 | 620 | 127 | 0.956 | 0.963 | 0.992 |
| s2_emb1 | 16502 | 504 | 72 | 0.966 | 0.970 | 0.996 |
| s2_emb3 | 20755 | 328 | 200 | 0.975 | 0.984 | 0.990 |

**Supplemental Table 2.** Cell Detection Performance on embGAN-processed DIC Images.

| embGAN | TP | FP | FN | Accuracy | Precision | Recall |
| --- | --- | --- | --- | --- | --- | --- |
| s1_emb2 | 21239 | 368 | 760 | 0.95 | 0.983 | 0.965 |
| s1_emb3 | 16113 | 187 | 673 | 0.95 | 0.989 | 0.960 |
| s2_emb1 | 18271 | 466 | 617 | 0.944 | 0.975 | 0.967 |
| s2_emb3 | 21358 | 467 | 773 | 0.945 | 0.979 | 0.965 |

**Supplemental Table 3.** Cell Tracking Performance on Fluorescence Images.

| Fluorescence | TP | FP | FN(missing) | FN(incorrect) | FN | Accuracy | Precision | Recall |
| --- | --- | --- | --- | --- | --- | --- | --- | --- |
| s1_emb2 | 20635 | 464 | 176 | 78 | 254 | 0.967 | 0.978 | 0.987 |
| s1_emb3 | 16407 | 640 | 128 | 48 | 176 | 0.953 | 0.962 | 0.989 |
| s2_emb1 | 16734 | 529 | 76 | 37 | 113 | 0.963 | 0.969 | 0.993 |
| s2_emb3 | 20935 | 390 | 270 | 200 | 470 | 0.961 | 0.982 | 0.978 |

**Supplemental Table 4.** Cell Tracking Performance on embGAN-Processed DIC Images.

| embGAN | TP | FP | FN(missing) | FN(incorrect) | FN | Accuracy | Precision | Recall |
| --- | --- | --- | --- | --- | --- | --- | --- | --- |
| s1_emb2 | 20693 | 730 | 766 | 440 | 1206 | 0.914 | 0.966 | 0.945 |
| s1_emb3 | 15830 | 416 | 682 | 294 | 976 | 0.920 | 0.974 | 0.942 |
| s2_emb1 | 17475 | 923 | 633 | 481 | 1114 | 0.896 | 0.945 | 0.940 |
| s2_emb3 | 20946 | 754 | 786 | 384 | 1170 | 0.916 | 0.95 | 0.947 |

## Methods

### *C*. *elegans* strains and culture

JIM113 was a gift of J. Murray (University of Pennsylvania) and N2 was sourced from the *Caenorhabditis* Genetics Center (University of Minnesota). Strains were grown at 20 ◻ on nematode growth media (US Biological) and fed OP50 *e. coli*. Gravid hermaphrodites from well-fed plates were cut using a needle in M9 buffer and embryos at the 2-cell or early 4-cell stage were mounted using a standard bead-mount approach. Briefly, extracted embryos were transferred via a hand drawn glass capillary to a 2 uL drop of M9 buffer containing ∽100 polystyrene beads 20 um in diameter (Polysciences Inc.), sandwiched between two pieces of #1.5 coverglass, and sealed with melted petrolatum jelly for imaging.

### DIC and fluorescence microscopy

Imaging was performed using either DIC alone for N2 or both DIC and fluorescence for JIM113 using an Olympus IX83 inverted frame equipped with a UPLSAPO60xs2 objective, a Visitech iSIM multipoint confocal scanner, ASI MX2000XYZ stage, and Hamamatsu Orca Fusion camera. The mCherry channel of JIM113 was acquired using 594 nm excitation and a 605 nm long-pass emission filter using 150 ms exposures and a laser power that was empirically tuned to not cause any qualitative developmental delays versus un-imaged control embryos and maintain a ∽100% hatch rate for imaged embryos. Embryos were imaged every 60 seconds with a 750 nm z spacing. DIC images were acquired with the Visitech scanner in brightfield bypass mode, a 50 ms camera exposure and the LED light source tuned to not generate any saturated pixels in the image. DIC illumination was generated using an Olympus UCD8 manual condenser equipped with a U525 oil immersion 1.4 NA top lens and a DICTHR tilt-shift slider. Images were acquired using micro-manager and cropped and converted to individual tiff volumes using Fiji. An additional dataset was acquired at a different site (California State University at Northridge) using a Leica DM6000 upright microscope equipped with an HCX PL APO 63x/1.4 NA objective, Leica K5 sCMOS camera, and DIC condenser equipped with a 1.4 NA oil immersion top lens. The DIC slider shear was adjusted empirically to approximate the contrast characteristics of images acquired on our Olympus microscope and images were acquired with a 50 ms exposure, 750 nm z-spacing, and illumination LED intensity adjusted to fill the sensor dynamic range without saturating pixels.

### embGAN

#### Image pre- and post-processing

Individual volumes were prepared for inference by the embGAN pipeline by performing per-volume contrast adjustment using Fiji’s built-in contrast adjustment pipeline set to target a maximum of 0.35% of pixels being saturated. The volumes were then converted to 8-bit grayscale and individual slices were exported as an image sequence. After inference, the 2D probability maps generated by embGAN were re-assembled into 3D stacks using a Fiji macro. Individual volumes were then clipped at an intensity of -4 by adding 4 to each image and setting all negative pixel values to 0.

#### embGAN implementation and training

embGAN is based on SeGAN, an adversarial generative model developed originally for the task of segmenting partially occluded objects in natural scenes. More specifically, embGAN is an adversarial training framework that utilizes a U-Net as a generator (Segmentor) and a multi-scale feature extractor (Critic) as input to a multi-scale objective loss function. This loss function is critical to the generalizable performance of embGAN as learning in adversarial networks often exhibits drastic instabilities.

#### Network Structure

The segmentor follows a typical U-Net structure i.e. a convolutional encoder-decoder with skip connections used to connect corresponding levels between the encoder and decoder. The encoder is composed of successive downsampling blocks, each followed by a residual block. The downsample block is comprised of a convolutional layer with stride of 2, each followed by a batchnorm layer and ReLU activation. The decoder mirrors the encoder and takes as input the output of the encoder. Each level of the decoder is made up of an upsampling block followed by a residual block. The upsampling block consists of a bilinear interpolation upsampling layer that performs upsampling by a factor of 2, a convolutional layer, batchnorm and ReLU activation. The residual block is used in both encoder and decoder and consists of a 1×1 convolution, a 3×3 convolution, followed by a 1×1 convolution. As in the U-Net, we add skip connections between corresponding layers of the encoder and decoder, concatenating the previous encoder outputs to the decoder output.

#### Critic

The critic performs feature extraction on the masked images, and is similar in structure to the encoder half of the segmentor without the residual blocks. It also makes use of global convolutions to increase the receptive field while reducing the number of learned parameters. Hierarchical features are computed at each level of C and are concatenated to produce the final output vector used as input to the multi-scale *I*_*MAE*_ loss.

In typical GAN frameworks, the critic network is trained to discern the difference between a ground truth or prediction generated via the segmentor network. To optimize model performance for segmentation we use the multiscale objective loss function L defined as.

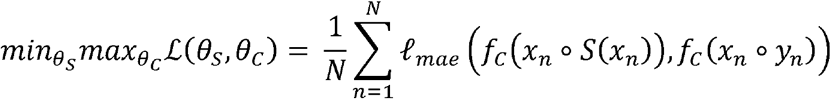

Where *I*_*MAE*_ is the Mean Absolute Error (MAE). The MAE is calculated at multiple scales n, between the input image masked by the predicted mask *x*_*n*_ *o S (x*_*n*_) and the input image masked by the ground truth labels. *x*_*n*_ *o y*_*n*_ . *f*_*C*_ represent the set of hierarchical features extracted by the critic network from each of the masked images. The *I*_*MAE*_ function is defined as:

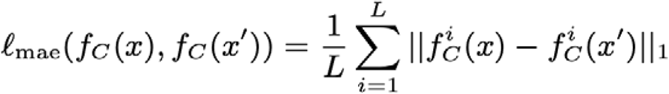

where L is the chosen number of scales in the critic network, and is the extracted feature map of image x at the ith layer of the critic network.

#### Label Generation

Labels were obtained from the fluorescence images using the pre-trained 2D_versatile_fluo model from Stardist.

#### Training

Both segmentor S and critic C networks are trained end to end in alternating fashion via backpropagation using a training set containing a total of 112,302 2D images. First, we fix S and train C doing one full forward and backward pass, then fix C and do the same for S for one step. The process of training S and C resembles a min-max game: S attempts to reduce the multi-scale feature loss, while C attempts to increase it. As the training progresses, both S and C networks improve their performance. Eventually, the segmentor can generate high quality predicted labels that closely match the ground truth masks. Both networks were trained simultaneously for 10,000 epochs with batch size 36 using the Adam optimizer and a learning rate of 0.0002.

### Automated lineage tracing and performance characterization

embGAN-processed images were processed using the latest release of StarryNite (https://github.com/zhirongbaolab/StarryNite) using a bifurcation classifier model trained using manually curated embGAN data and tuned parameter file (provided in our Github repository: https://github.com/shahlab-ucla/embGAN). For performance characterization, 4 embryos that were imaged separately from the original training set and not included in the training of embGAN were processed both using the fluorescence channel and a parameter file where only the intensity thresholds were adjusted and using embGAN-processed images as input. Curation was performed using the latest release of AceTree (https://github.com/zhirongbaolab/AceTree). All detections in the raw StarryNite output that were not linked to named cells in the curated version were counted as false positives (FP). Missing cells that were added manually during curation and counted as false negatives (FN). Errors in tracking were counted as FP if a nucleus was connected to an incorrect successor. We also defined two instances of FN for tracking: FN’s due to a missing detection, and FN’s due to a connection that was missed during the tracking stage. For cases with reciprocal errors, such as when tracks swap between two cells, previously described as “identity swap” errors, all erroneous edges are counted. In the simple case of two cells with swapped identities, a total of 4 errors are counted: two false negatives (FN) for the missing correct edge between each cell and the next time point, and two false positives (FP) for the incorrect edges passing between the tracks.

Tracing the lineages of the amphid neurons was done using AceTree with the embGAN-processed images of JIM113 embryos used in the performance characterization above except that only the sub-lineages that generate the two pairs of 12 amphid neurons on both left and right sides of the animal were edited. Errors were counted using a MATLAB script that compared the pointers in the StarryNite output files connecting detected cells at each time point between the edited and unedited files and every disagreement was counted as an error. All amphid neurons except for ASE, ASI, ASJ and ASK were tracked until the birth of the terminal neuron. These four were tracked until the birth of the neuroblast one cell cycle before the terminal neuron as the terminal neurons are born after an additional round of cell division compared to the remaining 8 amphid neurons. This terminal division often occurs after the beginning of embryonic twitching and thus was excluded from tracking in all embryos for performance characterization.

### Branch Distance Comparison of N2 and JIM113 Lineages

Whole-embryo and lineage-specific distances were computed using the intersection branch distance, defined as the *L*^*2*^ norm between a pair of ordered vectors containing the cell cycle times of the corresponding cells that exist in both lineages under comparison. Comparisons between the global clock of development between embryos were performed using the first principal component as a non-parametric measurement of the slope. Comparisons between intra-strain variability based on the median branch distance was performed using the rank sum test in MATLAB r2022a.

## Author Contributions

Conceptualization: PKS, MW, and ST. Software: MW, ST and GN. Validation and data analysis: RLJ, MW, AS, GN, and PKS. Supervision: PKS. Writing, original draft: PKS and MW. Writing, review, and editing: PKS, MW, RLJ, GN, and RLH. Funding acquisition: PKS and RLH.

## Acknowledgements

The authors would like to thank Neil Lin and Cho-Jui Hsieh for helpful discussions on style transfer robustness and generalization, and Alvaro Sagasti for encouragement and feedback. Some strains were provided by the CGC, which is funded by NIH Office of Research Infrastructure Programs (P40 OD010440). This work was supported by funding from NIH grant R21DC019485 to PKS.

